# mTORC1 Signaling Inhibition Modulates Mitochondrial Function in Frataxin Deficiency

**DOI:** 10.1101/2024.08.06.606942

**Authors:** Madison Lehmer, Roberto Zoncu

## Abstract

Lysosomes regulate mitochondrial function through multiple mechanisms including the master regulator, mechanistic Target of Rapamycin Complex 1 (mTORC1) protein kinase, which is activated at the lysosomal membrane by nutrient, growth factor and energy signals. mTORC1 promotes mitochondrial protein composition changes, respiratory capacity, and dynamics, though the full range of mitochondrial-regulating functions of this protein kinase remain undetermined. We find that acute chemical modulation of mTORC1 signaling decreased mitochondrial oxygen consumption, increased mitochondrial membrane potential and reduced susceptibility to stress-induced mitophagy. In cellular models of Friedreich’s Ataxia (FA), where loss of the Frataxin (FXN) protein suppresses Fe-S cluster synthesis and mitochondrial respiration, the changes induced by mTORC1 inhibitors lead to improved cell survival. Proteomic-based profiling uncover compositional changes that could underlie mTORC1-dependent modulation of FXN-deficient mitochondria. These studies highlight mTORC1 signaling as a regulator of mitochondrial composition and function, prompting further evaluation of this pathway in the context of mitochondrial disease.

## Introduction

Lysosomes and mitochondria must coordinate their highly specialized metabolic functions to help maintain cellular homeostasis under ever-changing nutrient and stressor conditions^1,2^. Lysosomes control mitochondrial composition, function and fitness via multiple mechanisms. The master growth regulatory kinase, mechanistic Target of Rapamycin Complex 1 (mTORC1) becomes activated at the lysosomal membrane in response to integrated signals from nutrients, growth factors, energy and oxygen^3,4^. From the lysosome, mTORC1 triggers pro-anabolic programs that, among other functions, reshape mitochondrial protein composition, respiratory capacity and fission-fusion dynamics^5,6^. Lysosomes also support mitophagy, the selective removal of portions of the mitochondrial network that become damaged upon metabolic, energetic or hypoxic stress, or due to aging^7^. Furthermore, lysosomes support mitochondrial function by controlling metabolite availability. and may engage in metabolite exchange though the establishment of contact sites that bring the membranes of the two organelles into close proximity^8,9,10,11^.

In periods of mitochondrial stress, mTORC1 activity can be inhibited either directly by AMPK or through regulatory control of its upstream modulators, GATOR1 or TSC1/2 complexes^12,13,14,15^. This inhibition is necessary to support upregulation of autophagy and catabolic pathways to reduce stress^13,16,17^. Accordingly, when lysosomes become dysfunctional, mitochondrial homeostasis is often impacted. In lysosomal storage disorders (LSDs), a family of rare diseases triggered by mutational inactivation of essential lysosomal enzymes and transporters, the primary lysosomal dysfunction invariably propagates to mitochondria^18^. In several LSDs, mTORC1 has been shown to be hyperactive; in one of these models, Niemann Pick Type C, inhibiting mTORC1 activity is sufficient to reverse some of the adverse mitochondrial effects, providing a mechanistic link between lysosomal and mitochondrial dysfunction^19,20^.

Consistent with the role of mTORC1 signaling in mitochondrial homeostasis, its pharmacological and genetic inhibition has proven beneficial in cellular and organismal models of mitochondrial diseases. For example, rapamycin treatment was shown to alleviate neurological symptoms in mouse models of Leigh Syndrome driven by defective assembly of mitochondrial complex I^21,22^. However, the molecular mechanisms underlying these effects remain unclear. More generally, the full range of transcriptional and post-transcriptional mechanisms via which mTORC1 controls mitochondria biogenesis and function in both normal and disease states remains to be fully elucidated.

Mitochondrial respiration and subsequent ATP production requires proper electron transfer and oxidation/reduction of various cofactors by the multi-subunit respiratory complexes I-IV, which constitute the electron transport chain (ETC)^23^. Accordingly, mutations in genes encoding for various core subunits of the respiratory complexes, or for their assembly factors and biosynthesis of cofactors, underlie a large family of inherited mitochondrial diseases. Among the most common mitochondrial diseases is Friedreich’s Ataxia (FA), an autosomal recessive condition caused by a triplet expansion in the gene encoding for the Frataxin (FXN) protein^24^. FXN is thought to function as an allosteric regulator for the synthesis of iron-sulfur (Fe-S) clusters, which are essential cofactors for the ETC^25,26,27^ .Thus, reduced FXN levels in FA lead to defective oxidative phosphorylation and energy metabolism, with severe impact in cardiac tissue and central nervous system among others.

In cellular and organismal models of FA, mTORC1 inhibition can extend lifespan and increase survivability^28,29^. Thus, Frataxin deficiency provides a useful model to interrogate by what mechanism(s) mTORC1 modulation may impact mitochondrial function and stress adaptation.

Here we sought to determine how chemical inhibition of mTORC1 impacts mitochondrial composition and function. We find that both allosteric and ATP-competitive inhibition of mTORC1 alter mitochondria respiration, polarization and mitophagy under stress. Focusing on cellular models of FXN depletion, here we uncover a role for mTORC1 inhibition in reprogramming mitochondria in a manner that may modulate their sensitivity to the loss of FXN protein.

## Results

### 1. mTORC1 inhibition impacts mitochondrial respiration and polarization

To establish whether mTORC1 modulation is influential on cell survival upon mitochondrial stress, we grew U2OS cells infected with shRNAs against either control LUC or DEPDC5, a component of the mTORC1-inhibiting GATOR1 complex^3,30^, to induce mTORC1 hyperactivation in U2OS cells^30^ (Fig. S1A). Indeed, when challenged by either the complex I inhibitor, Rotenone, or the complex V inhibitor, Oligomycin, to induce mitochondrial stress, DEPDC5-defective cells triggered a larger apoptotic response compared to control cells, as measured by fluorescence of a probe activated by caspase 3 or 7-dependent cleavage (Fig. 1A). Thus, mTORC1 hyperactivation appears to synergize with ETC-inhibiting compounds in disrupting cell viability.

**Figure 1.**
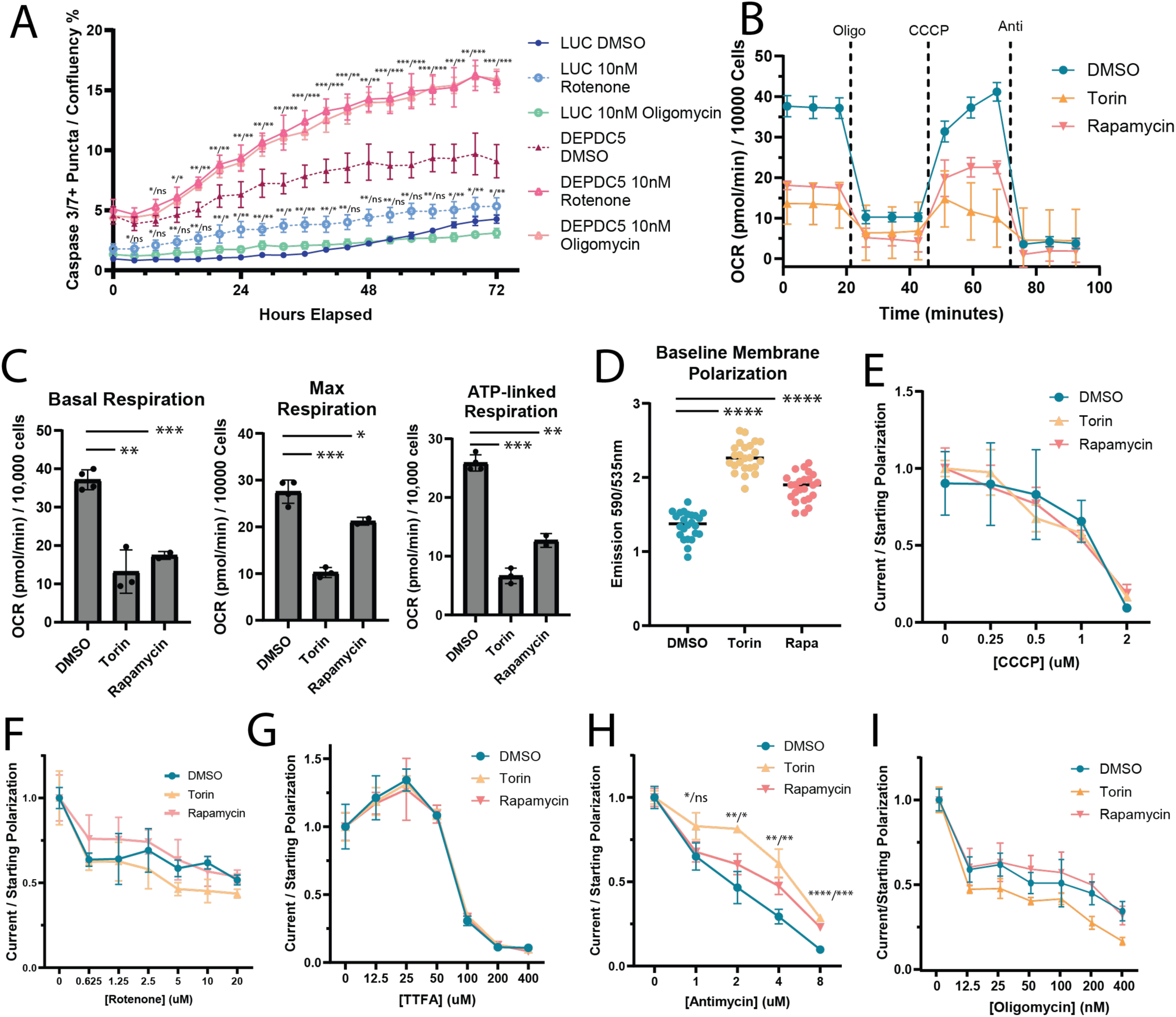
mTORC1 activity affects mitochondrial polarization and survivability to stress. **(A)** Timecourse for Caspase3/7+ puncta accumulation in U2OS cells infected with shRNAs for either DEPDC5 or LUC control that were treated with DMSO, 10nM Rotenone, or 10nM Oligomycin and measured for 3 days. P-values are marked where significance occurs and should be read between vehicle and drug conditions for shLUC (top), shDEPDC5 (middle), and shNPRL2 (bottom); **(B)** Oxygen consumption rate of DMSO, Torin, or Rapamycin treated U2OS cells; **(C)** Respiratory parameters calculated from Seahorse assay for cells inhibited with mTORC1 inhibitors; **(D)** Baseline mitochondrial membrane potential measured by JC-10 in either WT U2OS cells treated with DMSO, Torin or Rapamycin; **(E-I)** Percentage of baseline mitochondrial membrane potential remaining after treatment with dose response curves of CCCP (E) or ETC inhibitors Rotenone (F), TTFA (G), Antimycin (H), or Oligomycin (I). P-values are labeled where significance occurs and are read as p-value Torin/ p-value Rapamycin.

Using pharmacological inhibitors of mTORC1, we next determined the impact of acute mTORC1 inhibition on mitochondrial function and cell survivability. Specifically, U2OS cells were treated with the allosteric mTORC1 inhibitor, rapamycin, or with the catalytic mTOR inhibitor, Torin1, at concentrations 100-fold above their respective IC50 for 16-20 hours, followed by measurements of mitochondrial respiratory flux or membrane potential. Torin1, and to a lesser degree Rapamycin, decreased basal, ATP-linked and maximal respiration compared to vehicle-treated cells (Fig. 1B and 1C). Conversely, Torin1, and to a lesser degree Rapamycin, increased baseline mitochondrial membrane potential (MMP), as measured by the ratiometric MMP-sensitive dye, JC-10 (Fig. 1D). Prolonged mTORC1 inhibition was required for these effects to manifest; when either Torin1 or rapamycin were added during the experiment and OCR measured for two hours (a sufficient treatment time to impart signaling changes), the subsequent respiratory measurements were indistinguishable between vehicle-treated and mTORC1-inhibited cells (Fig. S1B).

The combination of increased MMP and decreased O_2_ consumption suggests that mTORC1 inhibition may differentially impact specific respiratory complexes. To examine this possibility, we measured to what extent mitochondrial membrane polarization is dissipated upon challenge with increasing concentrations of respiratory complex inhibitors and uncouplers, when mTORC1 is inhibited by Torin1 or Rapamycin. As a control, we used the proton uncoupler, Carbonyl cyanide m-chlorophenylhydrazone (CCCP), whose mechanism of action does not involve targeting a complex. Although the resting MMP was higher in mTORC1-inhibited versus control cells, CCCP led to a relative concentration-dependent drop in MMP that was indistinguishable between the two groups, as measured by JC-10 (Fig. 1E). Similar to CCCP, the complex-I inhibitor, rotenone, the complex-II inhibitor Thenoyltrifluoroacetone (TTFA), and the complex V (F-ATPase) inhibitor oligomycin showed identical dose-dependent alteration of MMP in control and mTORC1-inhibited cells (Fig. 1F-1G, 1I). In contrast Torin1, and to a lesser degree Rapamycin, antagonized dose-dependent MMP dissipation by challenge with the complex-III inhibitor, antimycin (Fig. 1H). Thus, mTORC1 inhibition suppresses mitochondrial respiration while increasing their baseline membrane potential, and counters dissipation of MMP triggered by treatment with a complex-III inhibitor.

Because loss of MMP potently stimulates mitophagy, and given the hyperpolarizing effect of mTORC1 inhibition, we evaluated whether Torin1 could affect the propensity of mitochondria to undergo depolarization-induced mitophagy. We incubated U2OS cells with CCCP to trigger PINK1-dependent phosphorylation of ubiquitin at Ser65, an event that results in subsequent capture of depolarized mitochondria by autophagic vesicles^31,32,33,34^. Accumulation of phospho-Ser65 ubiquitin (pS65-Ub) was significantly reduced in cells treated with Torin1 overnight compared to vehicle-treated cells (Fig. S1C). Because mTORC1 inhibition induces autophagy, we treated cells with the lysosomal deacidification agent Bafilomycin A1 (BafA1) after Torin1 treatment but before CCCP to prevent degradation of mitochondria captured via autophagy. This treatment did not restore pS65-Ub levels to those of cells treated with CCCP alone, indicating that the reduced levels of pS65-Ub in CCCP and Torin1-treated cells do not reflect increased autophagic flux (Fig. S1C).

### 2. Establishment and characterization of a cellular model of FXN depletion

We next asked how the mitochondrial functional parameters found to be influenced by mTORC1 behave in a mitochondrial disease model. To answer this, we use a cellular model of FXN loss. Because FXN is required for cell growth *in vitro* and thus cannot be knocked out completely^27,35^, we employed shRNAs in the U2OS cell line to establish a model in which FXN is reduced acutely (Fig. S2A). The inducible nature of this model allowed us to growth sufficient cells for both functional assays and biochemical fractionation, prior to acute FXN knock-down (Fig. S2A). This system stimulated the induction of energetic stress markers that signal mitochondrial dysfunction, such as ATF, SESN2, and phosphorylation of the AMPK target Acetyl-CoA Carboxylase (Fig. S2A and S2B). To independently corroborate key findings in this model, we also employed two FRDA patient-derived skin fibroblastic lines, along with a control fibroblastic line from a healthy individual (Fig. S2C).

FXN-depleted U2OS cells displayed multiple hallmarks of mitochondrial dysfunction relative to control (shLUC) cells. These knockdown cells showed reduced MMP at baseline (Fig. 2A) and increased apoptotic cell death following knock-down of FXN (Fig. 2B, Fig S2D). .Oxygen consumption rates in this acute knockdown model were not as severely diminished as in the FRDA patient fibroblasts, presumably due to the long stability of ETC component proteins^36^ (Fig. 2C and 2D). To characterize the potential ETC defect of FXN-depleted cells in more detail, we challenged them with increasing concentrations of uncouplers and respiratory complex inhibitors, and measured the resulting drop in MMP by JC-1 0. CCCP, rotenone and oligomycin dissipated the membrane potential of control and FXN-depleted U2OS cells with equal potency (Fig. 2E, 2F, 2I). In contrast, FXN-depleted cells enhanced the effects of both TTFA and antimycin on MMP (Fig. 2G, 2H). Thus, FXN loss decreased mitochondrial respiration, baseline polarization and maintenance of MMP upon both complex II- and complex-III inhibition.

**Figure 2.**
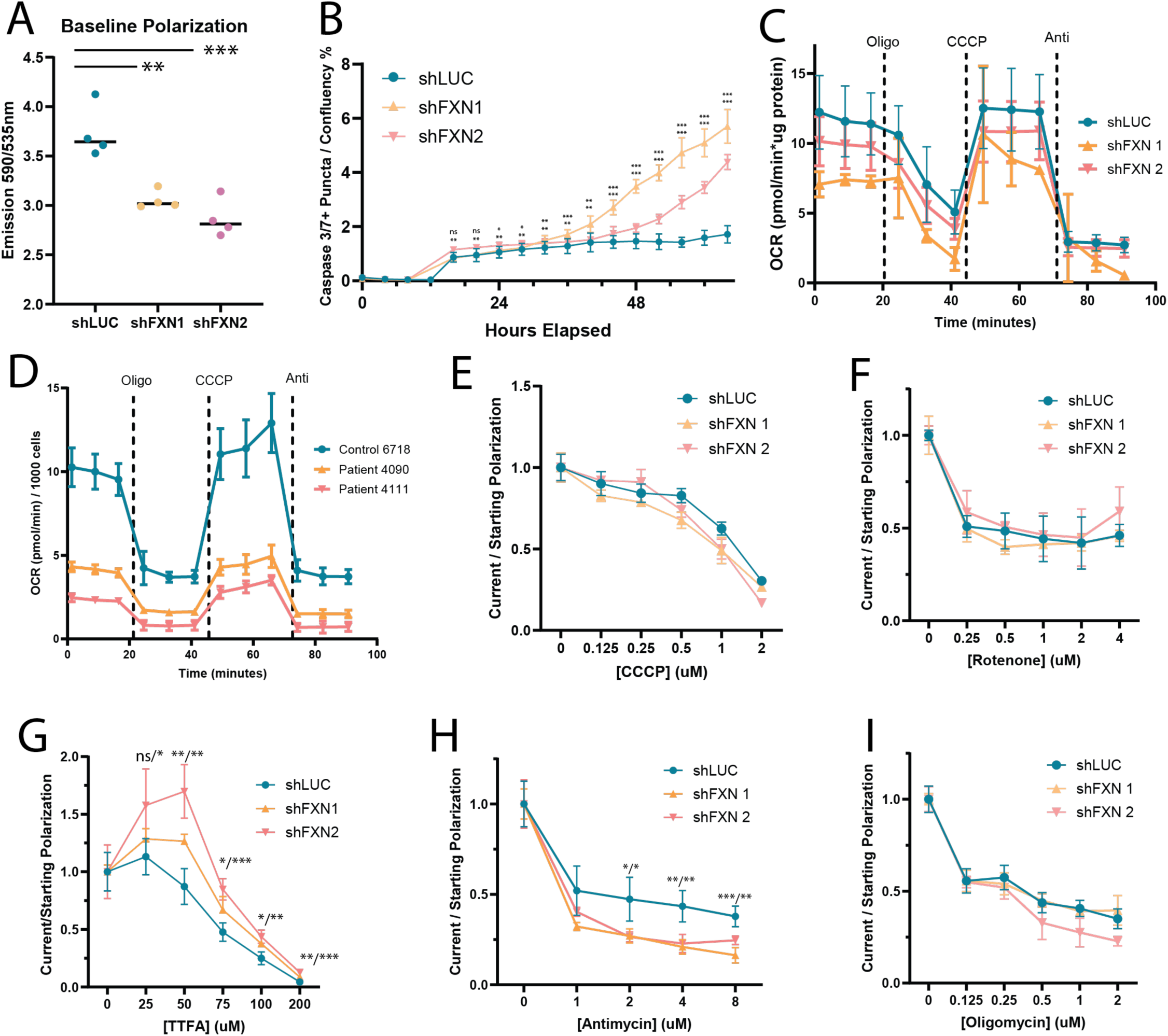
FXN-deficient cells have altered mitochondrial function. **(A)** Baseline mitochondrial membrane potential measured by JC-10 in shLUC or shFXN U2OS. P-value is <0.01 for shFXN1 and <0.001 for shFXN2; **(B)** Accumulation of caspase3/7-activated fluorescent probe in U2OS cells growing after infection with either shLUC, shFXN1, or shFXN2. P-values are marked where significant and should be read between the LUC and FXN hairpins, with FXN1 (top) and FXN2 (bottom); **(C, D)** OCR as measured by Seahorse in either shFXN U2OS **(C)** or FRDA patient **(D)** cells. **(E-I)** Percentage of starting MMP in shLUC or shFXN cells treated with varying concentrations of CCCP (E), Rotenone (F), TTFA (G), Antimycin (H), or Oligomycin (I). P-values are labeled where significance occurs and are to be read as p-value shFXN1/p-value shFXN2.

### 3. mTORC1 inhibition modulates mitochondrial function and survival of FXN-defective cells

We next determined how mTORC1 inhibition affects functional parameters of FXN-depleted mitochondria. Although FXN deficiency triggered mitochondrial stress responses, mTORC1 signaling was still active (FigS2A-C). As seen in control cells, treating FXN-depleted U2OS cells with Torin1 and Rapamycin increased their MMP (Fig. 3A). Moreover, both Torin1 and Rapamycin boosted MMP maintenance under increasing concentrations of antimycin in both control and FXN-depleted cells (Fig. 3B).

**Figure 3.**
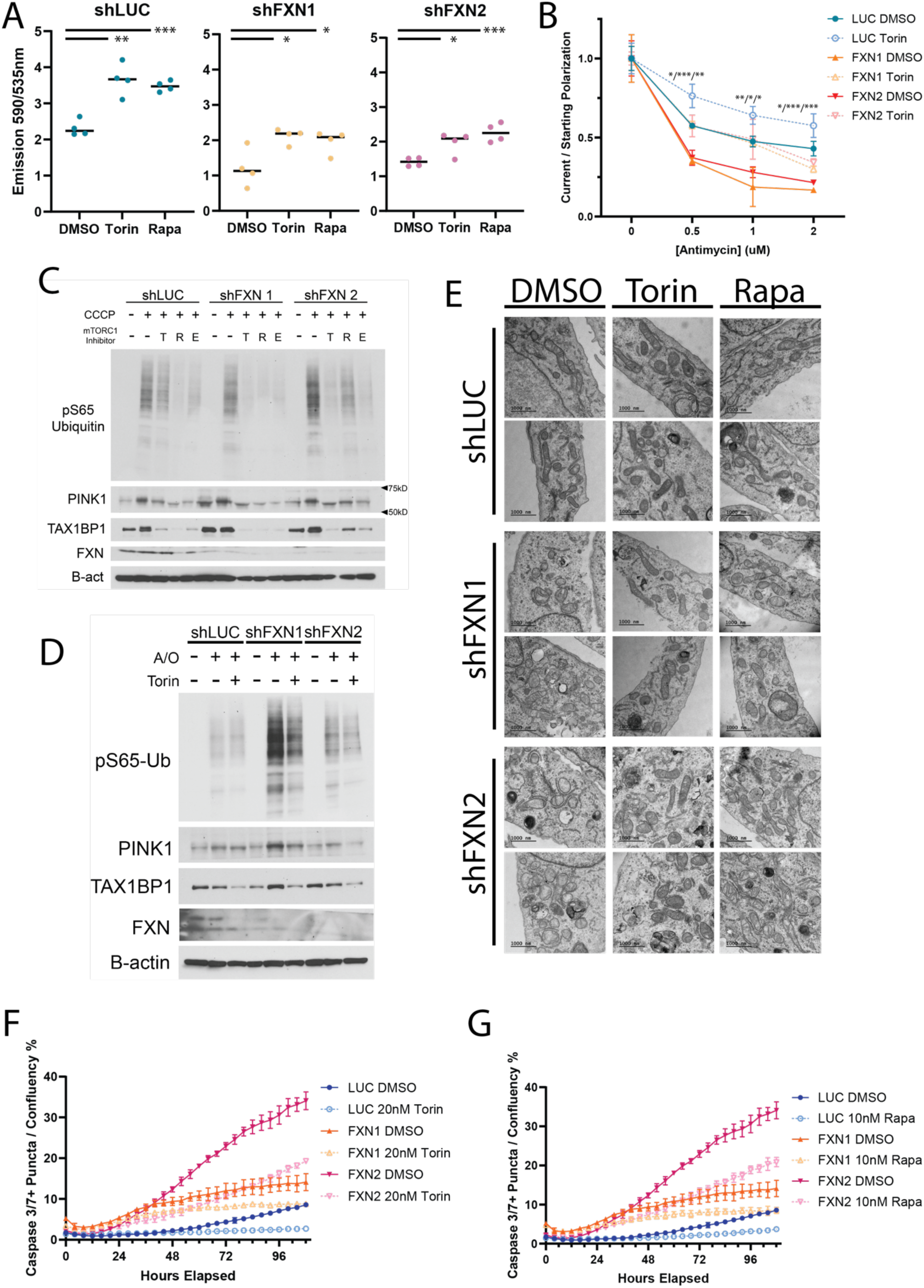
mTORC1 inhibition can offset some deficiencies of FXN-deficient mitochondria. **(A)** Membrane polarization of shLUC or shFXN U2OS cells treated with either DMSO, TorinA1, or Rapamycin; **(B)** Percentage of MMP loss incurred during Antimycin treatment in either shLUC or shFXN U2OS cells that have been treated with DMSO or TorinA1. P-values displayed signify significance between Torin and DMSO treated cells within the same genetic condition and are labeled where significance occurs; **(C,D)** Immunoblots for mitophagy markers in shLUC or shFXN cells that have been treated with DMSO or mTORC1 inhibitors overnight and then depolarized with CCCP (C) or Antimycin/Oligomycin (D); **(E)** Electron microscopy images of U2OS cells infected with either shLUC or shFXN and then treated with DMSO, TorinA1, or Rapamycin overnight; **(F)** Analysis of the electron density in mitochondria from EM images. For each mitochondrial particle, the integrated pixel value was divided by the particle area to establish the electron density measurement; **(E,F)** Caspase3/7+ probe activation in shLUC or shFXN U2OS cells grown in the presence of 20nM Torin or 10nM Rapamycin

Consistent with their reduced ability to maintain MMP, FXN-depleted cells showed enhanced accumulation of both PINK1 and p65Ub upon challenge with either CCCP or a combination of antimycin and oligomycin (the latter to prevent complex V from repolarizing mitochondria by working in reverse^37^) (Fig. 3C and 3D). Notably, pre-treating both control and FXN-depleted cells with Torin1, Rapamycin as well as the covalent mTORC1 inhibitor, EN-6^38^, strongly suppressed the accumulation of PINK1 and p65Ub triggered by either antimycin or CCCP (Fig. 3C and 3D). The same protective effects against depolarization-induced mitophagy were seen in the patient-derived fibroblasts (Fig. S2E).

The respective effects of FXN loss and mTORC1 inhibition on mitochondrial internal structure were analyzed via transmission electron microscopy (TEM). Here, control and FXN-depleted cells were treated with either vehicle, Torin1, or Rapamycin. Consistent with previous observations, relative to control, FXN-depleted mitochondria exhibited a loss of internal organization, as shown by the absence of visible cristae^39^ (Fig. 3E).

This loss of internal structure is a hallmark of swollen mitochondria, and can be quantified by the decreased electron density of the matrix^40^ (Fig. S2F). Neither Rapamycin nor Torin1 appeared able to restore the morphology of the FXN-depleted mitochondria to healthy-like: in mTORC1-inhibited, FXN-depleted cells, mitochondria remained generally fragmented and internally disorganized. However Torin1, and to a lesser degree rapamycin, caused the mitochondria matrix to acquire a more condensed, electron-dense appearance irrespective of FXN status, a phenotype associated with decreased oxidative phosphorylation (Fig. 3E)^41,42^.

Loss of mitochondrial internal organization is associated with release of cytochrome C, leading to apoptotic cell death. Consistently, whereas shRNA-mediated FXN depletion was sufficient to induce apoptosis (Fig. 2D), maintaining FXN-depleted cells in rapamycin or Torin1 reduced accumulation of the cleaved caspase 3/7 marker (Fig. 3F and 3G).

Collectively, these results indicate that the previously established effects of mTORC1 inhibition, namely enhanced maintenance of MMP, resistance of complex-III to challenge by antimycin, and reduced depolarization-dependent p65Ub accumulation, also occur in FXN-depleted cells. These changes co-occur with, and may be linked to, the decreased susceptibility of mTORC1-inhibited cells to apoptotic cell death triggered by FXN depletion.

### 4. mTORC1 inhibition reprograms the mitochondrial proteome in FXN-defective cells

To gain insights into the mechanisms of mTORC1-dependent modulation of mitochondrial function, we carried out mitochondria immunoisolation and proteomic profiling from FXN-depleted cells^43^. The HA^x3^-GFP-OMP25 mitochondria affinity tag was introduced in U2OS cells that were subsequently depleted for FXN via shRNA. These lines were treated with vehicle or Torin1 overnight, followed by cell fractionation and rapid mitochondria pull-down (Fig. 4A, S3A). Label-free quantitation (LFQ) was then used to determine mitochondrial proteome changes due to loss of FXN, as well as the effect of Torin1 in each background (Supplementary data 1). We focused our analysis to proteins identified as mitochondrial through the MitoCarta 3.0 database^44^.

**Figure 4.**
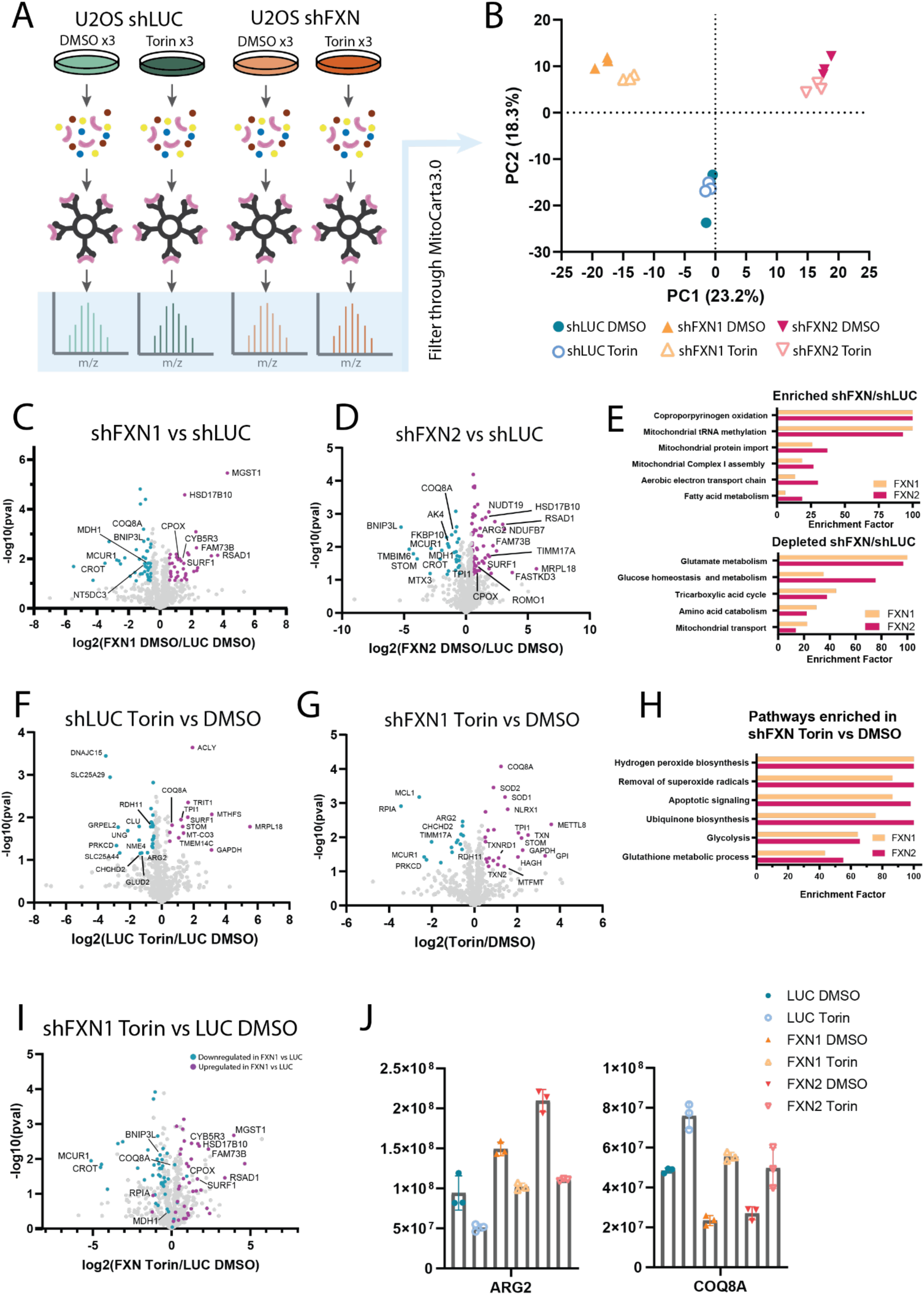
Mitochondrial proteomics in Torin treated, FXN-deficient cells identify machinery that may be involved in mTORC1-mediated mitochondrial function. **(A)** Schematic of mitoIP protocol; **(B)** Principle-component analysis of the mitochondrial proteome in each sample type; **(C and D)** Volcano plots displaying changes in mitochondrial genes in vehicle-treated cells comparing FXN-deficient with wildtype. Significantly altered genes (pval =<0.05) highlighted in blue (>40% decrease of protein levels) or fuschia (>40% increase of protein levels); **(E)** Enrichment analysis for pathways altered during Torin deficiency in both hairpins. **(F and G)** Volcano plots displaying the alteration of mitochondrial proteome upon mTORC1 inhibition by Torin1 in control shLUC (F) and shFXN (G) cells. **(H)** Pathway analysis for proteins enriched in mTORC1-inhibited FXN-deficient cells; **(I)** Volcano plot displaying changes in the mitochondrial proteome between Torin1-treated, FXN-deficient cells and vehicle-treated control shLUC cells. Highlighted proteins are the same as those highlighted in (C) to display how mTORC1 inhibition by Torin1 affects the machinery altered by FXN deficiency. **(J)** Bar plots of LFQ values for genes identified to restore to control baseline upon Torin1 treatment in FXN-deficient cells.

Principal component analysis (PCA) showed that FXN-depleted mitochondria were clearly separated from control mitochondria (Fig. 4B). However, the two shFXN samples were also separate from each other, possibly reflecting different degree of FXN knock-down, as well as noise resulting from sample preparation. Despite differences in individual protein levels, many pathways overlapped between the two FXN-depleted samples compared to shLUC-transduced samples. Specifically, mitochondria purified from cells transduced with either FXN-targeting shRNAs displayed an increase of proteins involved in heme metabolism, such as coproporphyrinogen oxidase (CPOX) and radical S-adenosylmethionine domain containing 1 (RSAD1), whereas proteins involved in glucose metabolism and the TCA cycle were significantly depleted (Fig. 4C-4E).

To gain insights into functional changes that may occur upon mTORC1 inhibition, we compared the Torin1-induced changes in both control and FXN-depleted mitochondria, relative to their vehicle-treated counterparts (Fig. 4F-4H and S3B-S3C). Across control and both FXN-depleted mitochondria samples, proteins involved in oxidative stress were similarly altered (Fig 4H and S3D). The superoxide dismutases 1 and 2 (SOD1 and SOD2), were enriched by Torin1 treatment, whereas coiled-coil-helix-coiled-coil-helix domain containing 2 (CHCHD2), a protein involved in hypoxic stress response, was depleted^45,46,47^ (Fig 4H and S3D).

Additionally, glycolytic proteins triosephosphate isomerase (TPI) and glyceraldehyde 3-phosphate dehydrogenase (GAPDH) increased upon Torin treatment. On top of their roles as glycolytic enzymes, both TPI and GAPDH have been found to alter mitochondrial redox stress^48,49^. Notably, in mitochondria from both FXN-depleted lines, but not control cells, Torin1 treatment led to increased levels of proteins involved in glutathione metabolism and ubiquinone biosynthesis, whereas these categories were not altered by Torin1 treatment in control cells (Fig. 4H and S3C).

Finally, we were able to identify a small number of proteins whose levels were altered in FXN-depleted versus control mitochondria, but which were restored to control levels by Torin1 treatment. Amongst this subset are Coenzyme Q8A (CoQ8A), a protein component of the CoQ biosynthesis cascade, as well as Arginase 2, which promotes oxidative activity through Complex II activation^50,51^ (Fig. 4I-4J, S3D-S3E).

These mitochondrial proteomic data begin to shed light on the changes in mitochondrial composition that occur as a function of FXN loss. They also point to compositional changes that could underlie the functional reprogramming mediated by mTORC1 inhibitors in both control and FXN-defective cells.

## Discussion

Here the effects of acute pharmacological inhibition of mTORC1 signaling on mitochondrial composition and function were examined both in healthy cells and following depletion of the essential Fe-S cluster biogenesis factor, Frataxin^52^. Blocking mTORC1 resulted in pronounced suppression of mitochondrial respiration, an effect in line with previous reports^6^ that was accompanied by increased polarization at baseline. These two effects are potentially linked as respiration consumes the proton gradient underlying MMP to power ATP generation; if confirmed, this would represent an mTORC1-dependent decoupling mechanism to promote MMP maintenance. In that respect, the apparent ability of mTORC1 inhibitors to counter MMP depletion caused by antimycin points to complex-III modulation as a potential mechanism. An alternative possibility is that mTORC1 inhibition may cause the F-ATPase to run in reverse, thereby consuming ATP to maintain MMP^37^. Furthermore, the ‘condensed’ appearance of the mitochondrial matrix in electron micrographs of Torin1- and rapamycin-treated cells further supported a generalized decreased in mitochondrial activity upon mTORC1 inhibition.

The effects of Torin1 and rapamycin were similar in healthy and FXN-depleted cells, in that mTORC1 inhibition decreased respiration and hyperpolarized both control and FXN-depleted mitochondria. Moreover, mTORC1 signaling appeared not to be altered by FXN depletion. Based on these criteria, mTORC1 would not quality as a modifier of FXN deficiency. However, cell death caused by FXN depletion was significantly reduced upon treating FXN-defective cells with mTORC1 inhibitors, indicating that one or more pathways downstream of mTORC1 inactivation can reduce the toxicity caused by FXN loss.

mTORC1 regulates both anabolic and catabolic pathways, and modulation of several of them, including translation initiation and autophagy could have profound effects on mitochondrial composition and function^4,5,6^. Interestingly, mTORC1 inhibition appeared to reduce mitophagy caused by mitochondrial depolarizing agents. A reason for this could be the reduced stabilization of PINK1 that was observed in cells (both control and FXN-depleted) that were co-treated with mitochondrial depolarizing agents and with Torin1 or rapamycin. In turn, the reduced PINK1 stabilization led to lower levels of Ser65-phosphorylated ubiquitin, which ultimately drives mitochondrial engulfment by autophagosomes and their delivery to lysosomes^53^. How mTORC1 inhibition leads to reduced PINK1 stabilization in depolarized mitochondria remains to be determined, although boosting of baseline MMP could be a factor.

Finally, comparing the proteomics of control and FXN-depleted mitochondria unveils several pathways that are modified by mTORC1 may contribute to counter mitochondrial stress induced by FXN loss, including oxidative stress response and cofactor synthesis.Within this pathway, CoQ8 was notable in that its reduced levels in FXN-depleted mitochondria were restored to control levels upon mTORC1 inhibition. CoQ plays a key role in the ETC by shuttling electrons to complex-III. Assuming that the increased levels of CoQ biosynthetic enzymes translates to higher CoQ_10_ levels in Torin1-treated mitochondria, the additional CoQ_10_ may compete with antimycin for binding to the quinone reduction site of complex-III. This could provide a mechanistic rationale for the increased resistance of mTORC1-inhibited mitochondria to antimycin-dependent depolarization but not to other ETC inhibitors. In turn, higher MMP may inhibit release of cytochrome C from stressed mitochondria, thereby explaining the reduced apoptotic rate of FXN-depleted cells upon mTORC1 inhibition.

In conclusion, while irrespective of FXN levels, the multi-pronged effects of chemical mTORC1 modulation on mitochondrial activity add up to a pro-survival effect on FXN-depleted cells. While these effects require further mechanistic investigation, their elucidation could point to pharmacological mTORC1 modulation in the context of FXN loss and other forms of mitochondrial dysfunction.

## Acknowledgements

We thank M Napierla at UT Southwestern for providing patient-derived FA cell lines, and A. Jain and R. Peruzzo for critical reading of the manuscript.

## Materials and Methods

### Mammalian Cell Culture

WT HEK293T and U2OS cells were cultured in DMEM (Gibco, 11965-092) supplemented with 10% (v/v) fetal bovine serum (Sigma, F0926), 100U/mL penicillin, and 100µg/mLstreptomycin (Gibco, 15140-122. FRDA patient lines were purchased from the Napierala Lab’s FRDA Cell Line Repository at the University of Texas Southwestern (Control: #6718, Patient 1: 4090, Patient 2: 4111)and cultured in Gibco DMEM supplemented with 15% (v/v) fetal bovine serum, 1X MEM Non-Essential Amino Acids Solution (Gibco, 11140076), 1mM Sodium Pyruvate (Gibco, 11360070), 100U/mL penicillin, and 100µg/mL streptomycin. Cell Lines used were labeled as follows: Control: All cells were cultured at 37C and 5% CO_2_.

### Cell Line Generation

A lentiviral vector encoding the cDNA for the MitoIP tag was generated as follows. The coding block for the 3xHA-eGFP-OMP25 tag was cloned from its original pLJM1 plasmid^43^ and inserted into the pLVX lentiviral vector with a hygromycin resistance marker.

Short hairpin oligonucleotides (shRNAs) were cloned into the pLKO.1 vector (The RNAi Consortium, Broad Institute). The shRNAs used in this study are as follows: FXN1 (TRCN0000377277), FXN2 (TRCN0000006137), DEPDC5 (TRCN0000134778), NPRL2 (TRCN0000234677), LAMTOR4 (TRCN0000281609), LAMTOR5 (TRCN0000153443), and LUCIFERASE non-targeting control (TRCN0000072243).

All lentiviral constructs were packaged into viral particles by co-transfection with psPAX2 (Addgene 12260) and pMD2G (Addgene 12259) constructs in HEK293T cells using PEI transfection reagent. Viral supernatant was captured at 48hr and again at 72hr, centrifuged to clear, and concentrated using Lenti-X Concentrator (Clontech, 631231) according to manufacturer’s protocol. Target cells were mixed with virus and 8µg/mL polybrene (Millipore, TR-1003-G) and then in a 6-well plate (stable generation) or their relevant assay dishes (knockdowns). After 24hr, the viral containing media was removed and replaced with selection of either 5µg/mL puromycin or 125µg/mL hygromycin.

### Drug Treatments

Unless otherwise stated, drug treatments were performed as follows. Rapamycin (Calbiochem, 553210) was used at 100nM for 20hr. TorinA1 (Tocris, 4247) was used at 250nM for 20hr. For membrane potential experiments, CCCP (MedChemExpress, HY-100941) was used for 20min, Rotenone (Calbiochem #557368) was used for 40min, Antimycin A (Sigma-Aldrich, A8674) was used for 45min, Oligomycin (Sigma-Aldrich O4876) was used for 15min, and 2-Thenoyltrifluoroacetone (TTFA) (Thomas Scientific, C818L50) was used for 3hr. For mitophagy induction, CCCP was used at 10µM for 5hr and Antimycin A and Oligomycin A were used at 10µM and 5µM, respectively, for 6hr.

### Cell growth and apoptosis

U2OS Cells were seeded at 3,000/well in a 96-well plate and left to adhere overnight. The following day, the seeding media was swapped out for fresh media that contained vehicle/drug and 2.5nM CellEvent Caspase3/7+ Green Probe (Invitrogen, C10432). Then, confluence and number of green puncta were assessed by IncucyteS3 (Sartorius) at 4hr intervals. The green puncta count was normalized to confluence percentage to account for changes in growth rate across cell lines and drug treatments.

### Mitochondrial Membrane Potential

Cells were seeded in a 96-well plate. Before the assay began, fresh media was added to each well. Then, depolarizing drugs (CCCP, Antimycin A, Rotenone, Oligomycin, or TTFA) were added to their final indicated concentrations for the above-described amount of time. Additionally, JC-10 (Enzo Life Sciences, 52305) was added to a final concentration of 5uM and incubated for 30min total. When the incubations had finished, cells were rinsed 1X with PBS and then another round of PBS was added. Then, the plate was analyzed by a TECAN Spark (Model#30086376) by fluorescent readout for green monomeric state (Ex. 485nM, Em. 535nM) or red aggregated polarized state (Ex. 535nM, Em. 590nM). Background signal in each channel was calculated from wells with cells that had not been treated with JC-10 dye and subtracted before the aggregate/monomer ratio was calculated. For drug curves, the percentage of starting polarization was calculated by determining the average ratio of each condition without depolarization agent and dividing each individual replicate’s value by this number.

### Seahorse Respiration Assay

Oxygen consumption rate was analyzed by a Seahorse XFe24 Analyzer (Agilent). Cells were seeded in culture media. An hour before the experiment, media was rinsed 1X with PBS and replaced with DMEM without sodium bicarbonate, pH 7.4 and equilibrated at 37C without supplemental CO_2_. OCR was then recorded using Agilent’s Mito Stress Protocol, involving treatment with 500nM Oligomycin, 2µM CCCP, and 1µM Antimycin. After the experiment, cells were rinsed 1X with PBS and fixed with 4% PFA for 20min before rinsing 1X more with PBS and staining with Hoescht in PBS (1µg/mL, 30min). Staining solution was once again rinsed and replaced with PBS and cell count was analyzed using a Biotek Cytation1 (Agilent) measuring puncta count at Ex.377, Em.447. The OCR for each well was then normalized to its corresponding cell count.

### Cell Lysis and Immunoblotting

Cells were rinsed 1X with PBS and then scraped into lysis buffer (40mM HEPES, 10mM sodium pyrophosphate, 10mM β-glycerol phosphate, 4mM EDTA, 1% TritonX-100, pH7.4, supplemented with Pierce protease and phosphatase inhibitor tablets (Pierce, A32963)) and incubated at 4°C with rocking for 1hr to ensure complete lysis. Lysates were then spun at 17,000 x g for 10min at 4C and the pellet was removed.

Protein concentrations were standardized by BCA analysis using a BCA Protein Assay Kit (Pierce, 23225) according to the manufacturer’s instruction. Samples of equal concentration were prepared for SDS-PAGE by addition of 5X SDS sample buffer (235nM Tris pH 6.8, 10% SDS, 25% glycerol, 25% β-mercaptoethanol, 0.1% bromophenol blue) and denatured for 10min at 95°C. 10µg of each sample was loaded on either a 12% (Invitrogen, XP00122) or 4-20% (Invitrogen, XP04205) Tris-glycine gel and resolved by electrophoresis in Tris-Glycine running buffer (25mMTris, 190-mM glycine, 0.1% (w/v) SDS). Proteins were transferred to a PDVF membrane (Millipore, IPVH00010) by wet transfer at 34V for 2hr10min in a CAPS transfer buffer (10mM CAPS, 10% EtOH, pH 11). Then, membranes were blocked in either 5% non-fat milk or 5% BSA in TBS-T and incubated overnight at 4C with primary antibodies (diluted in either 5% non-fat milk or 5% BSA in TBS-T). Phosphorylation-site antibody dilutions used PBS-T in place of TBS-T. Membranes were rinsed with TBS-T and incubated with horseradish peroxidase-conjugated secondary antibodies diluted in 5% non-fat milk, TBS-T for 1hr at room temperature. Membranes were again rinsed with TBS-T and incubated with ECL Blotting Substrate (Pierce 32106) before exposure to ProSignal Blotting Film (Prometheus 30-810L). Immunoblots of mitoIP samples were conducted in the same manner, instead using 2ug of sample and activated with Clarity Western ECL Substrate (BioRad, 170-5061).

### Antibodies

4EBP1 (Cell Signaling Technology, 9644S); pS65-4EBP1 (Cell Signaling Technology, 13443S); Acetyl-CoA Carboxylase (Cell Signaling Technology, C83B10); pS79-Acetyl-CoA Carboxylase (Cell Signaling Technology, 3661S); AMPK (Cell Signaling Technology, 5831T); pT172-AMPK(Cell Signaling Technology, 2535S); β-actin (Cell Signaling Technology, 8457S); DEPDC5 (Abcam, ab213161); FXN (Proteintech, 14147-1-AP); LC3B (Cell Signaling Technology, 3838S); p62 (Cell Signaling Technology, 39749S); PARP (Cell Signaling Technology, 9542S); PINK1 (Cell Signaling Technology, 6946S); S6K (Cell Signaling Technology, 2708S); pT389-S6K (Cell Signaling Technology, 9234S); SDHB (Proteintech, 10620-1-AP); SESN2 (Cell Signaling Technology, 8487S); TAX1BP1 (Cell Signaling Technology, 5105S); TOM40 (Santa Cruz Biotechnology, sc-11414); α-tubulin (Cell Signaling Technology, 2125S); pS65-Ubiquitin (Cell Signaling Technology, 70973S); VDAC (Cell Signaling Technology, 4661S)

### Transmission Electron Microscopy

After growth and treatment, cells were fixed with a 2% glutaraldehyde solution at 37C for 30min. Cells were then scraped and collected, rinsed 3X with PBS, and added to a solution of 1% Osmium Tetroxide + 1.6% Potassium ferricyanide in PBS for 1-2hr. Cells were rinsed 3X with PBS then 1X with distilled water before dehydration with acetone, using increasing concentrations from 35% upwards until they were fully dehydrated with 100% acetone, incubating for 10 min at each step. Samples were then infiltrated with 2:1 Acetone: (resin + BDMA) for 1hr, 1:1 for 1hr, and then 1:2 overnight. The next day they were incubated with 100% resin 3X for 1-2hr at a time, then embedded and baked at 60°C for 2days. When finished, the samples were imaged using a TECNAI 12 TEM.

### RT-qPCR

mRNA was isolated from cells using the BioRad Aurum™ Total RNA Mini Kit (BioRad, 7326820). RNA concentrations were assessed using a Nanodrop reading absorbance at 260nm. 1ug of RNA from each sample was then used for cDNA synthesis using the iScript™ Advanced cDNA Synthesis Kit for RT-qPCR (Bio-Rad). Real-time quantitative PCR was then performed in quadruplicate using a [check model] and detected using SYBR Green (Bio-Rad, 172-5271). The resulting CT values from melting curves were then normalized to those of a reference gene to calculate fold change.

#### Primers used were as follows

HPRT (F – TGACACTGGCAAAACAATGCA, R – GGTCCTTTTCACCAGCAAGCT); β-actin (F – ATTGCCGACAGGATGCAGAA, R – ACATCTGCTGGAAGGTGGACAG), ATF4 (F - CTTCACCTTCTTA CAACCTCTTC, R – GTAGTCTGGCTTCCTATCTCC), SESN2 (F – AGATGGAGAGCCGCTTTGAGCT, R – CCGAGTGAAGTCCTCATATCCG), FXN (F – ATCTTCTCCATCCAGTGGACCT, R – GCTGGGCATCAAGCATCTTTT)

### Mitochondrial Immunoprecipitation

Mitochondria were isolated from confluent 15cm dishes of U2OS cells by the mito-IP protocol established by^43^, with the modifications that cells were lysed with a syringe and 27G needle instead of a dounce homogenizer and samples were incubated with beads for 20min. The captured mitochondria were released by incubating the beads at 37°C for 1hr in KPBS + 0.1% NP-40 and the resulting protein normalized by BCA for western blotting, or in PBS + 0.05% ProteaseMax Surfactant (Promega V2071) and snap-frozen in liquid nitrogen for mass spectrometry analysis. For proteomics samples, 10% of the resulting elution volume was withheld and boiled with 5% SDS sample buffer to assess mitochondrial enrichment by western blotting.

### Proteomics Analysis

MitoIP proteomics samples were prepared in triplicate. Samples were analyzed by label-free qualitative LC/MS-MS using a Proxeon EASY-nanoLC system (ThermoFisher) coupled to a Orbitrap Fusion Lumos Tribid mass spectrometer (Thermo Fisher Scientific). The resulting mass spectra were analyzed with MassQuant software version 1.6.11.0 and filtered for contaminants from the GPM cRAP Sequences database. The resulting iBAQ values for these genes were normalized to total BAQ sum to calculate the label free quantification (LFQ) values for comparison. These LFQ values were then processed in R as follows: Proteins were filtered out if they had sequence coverage <5%. The average background value of each gene (calculated from the “blank” sample) was removed from each sample’s LFQ value. Any resulting value of 0 was replaced with a randomly generated number matching the bottom 1% of LFQ values to allow for statistical calculations. Statistical significance of proteins between samples was established by a parametric t-test.

---

**Figure S1.**
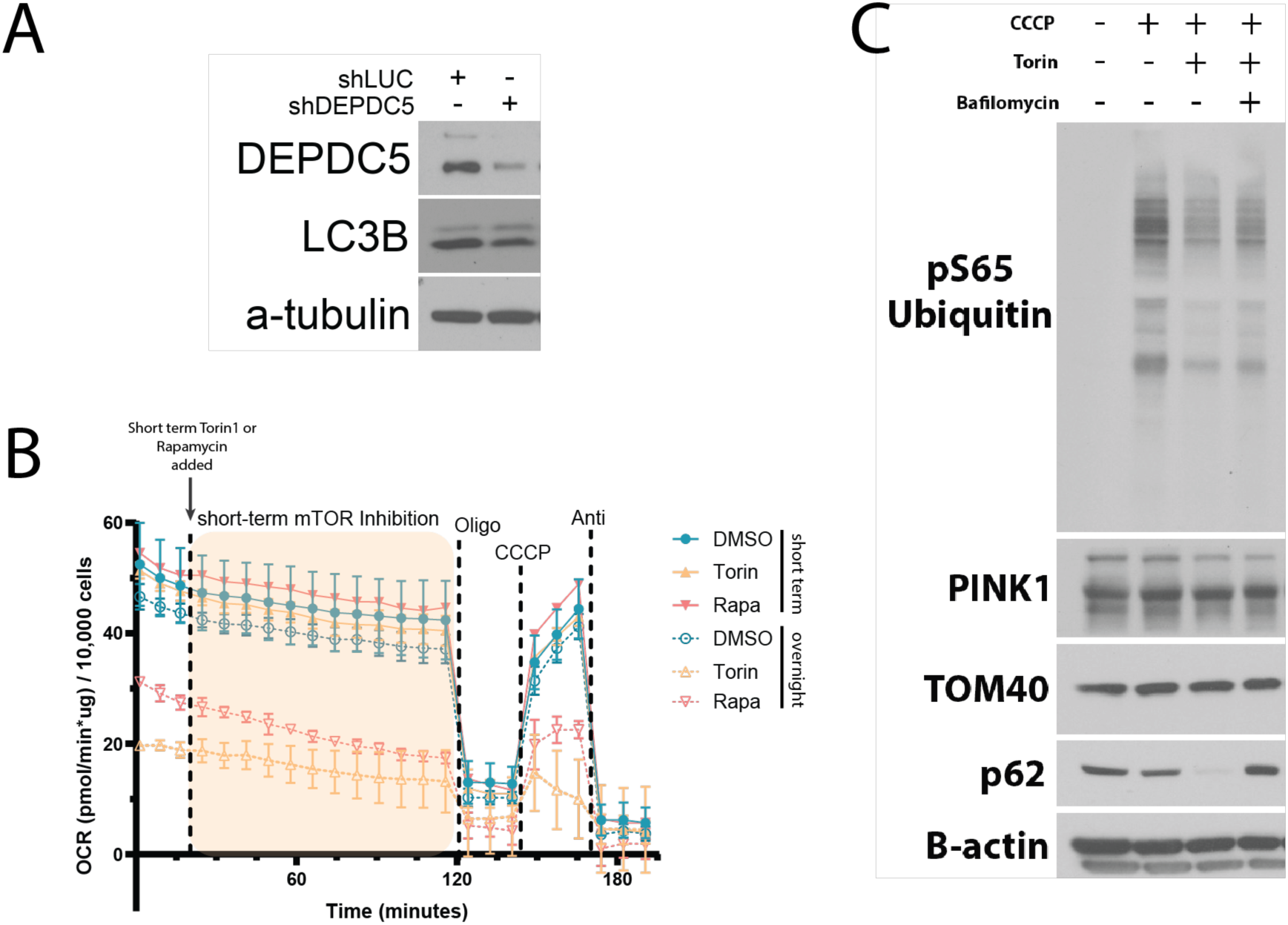
Affects of mTORC1 inhibition that influence mitochondrial function are the result of a longer term inhibition rather than immediate signaling effect. **(A)** DEPDC5 shRNA test for Fig1A control; **(B)** Western blot for mitophagic and mTORC1 signaling markers after either long- or short-term Torin1 treatment followed by CCCP challenge; **(C)** Western blot for mitophagic markers after treatment with Torin1 for 20hr, BafilomycinA1 for 12hr, and CCCP for 5hr; **(D)** OCR analysis of U2OS cells treated with vehicle, Torin1, or Rapamycin either overnight or for 1.5hr after taking basal measurements.

**Figure S2.**
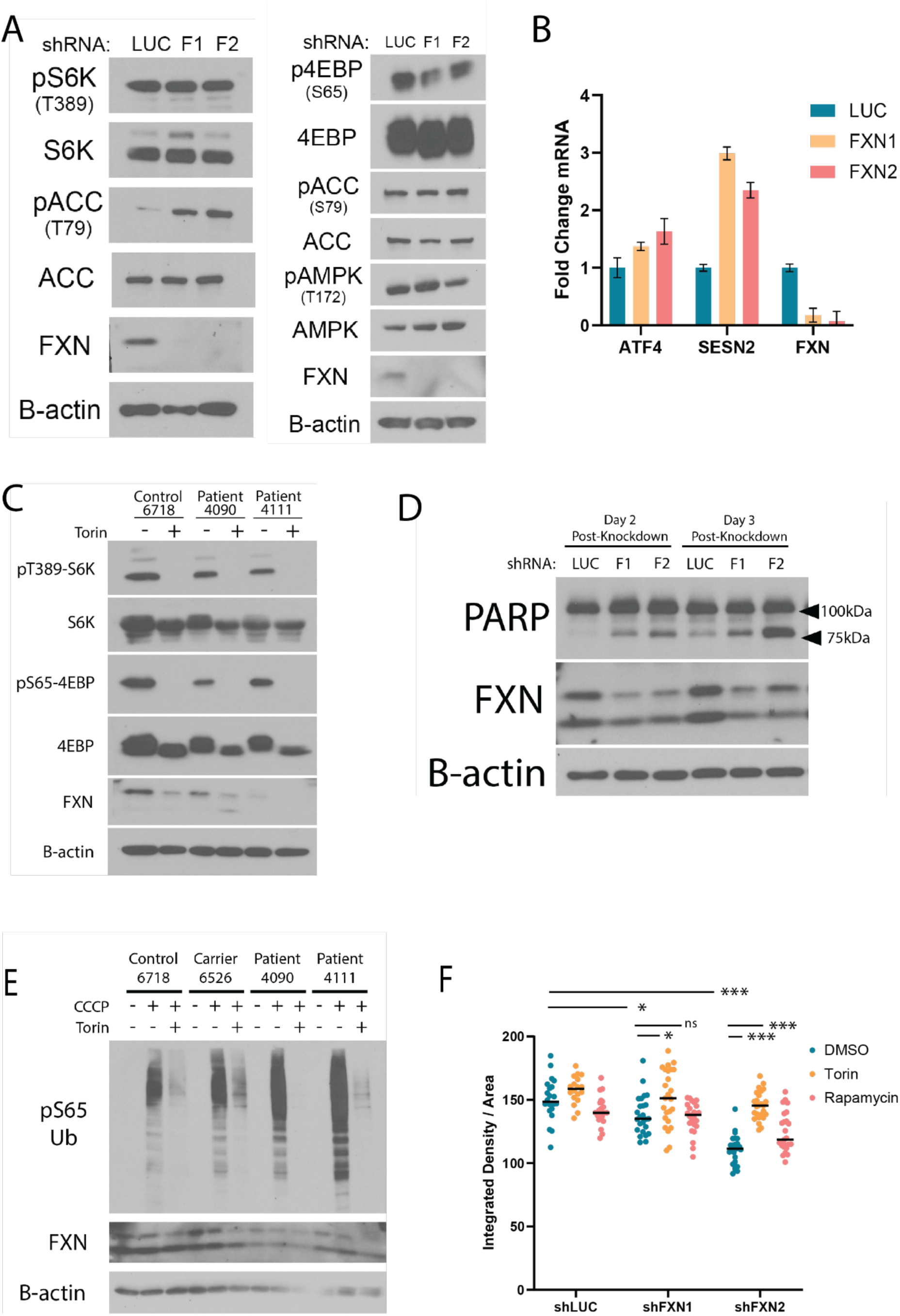
Model of FXN deficiency by shRNA knockdown induces mitochondrial stress markers without inhibition of mTORC1. **(A)** Western blot for mTORC1 and mitochondrial stress signaling markers in shLUC or shFXN cells; **(B)** qPCR for ATF4 and SESN2 activation in shLUC or shFXN cells. **(C)** Western blot of mTORC1 signaling markers in patient FA fibroblasts; **(D)** Western blot for cleaved PARP accumulation for two- and three-days post-shRNA mediated knockdown in U2OS cells; **(E)** Western blot of mitophagic markers in FRDA patient lines treated with Torin1 overnight and then challenged with CCCP; **(F)** Analysis of Integrated density/Area for mitochondria from Fig3E.

**Figure S3.**
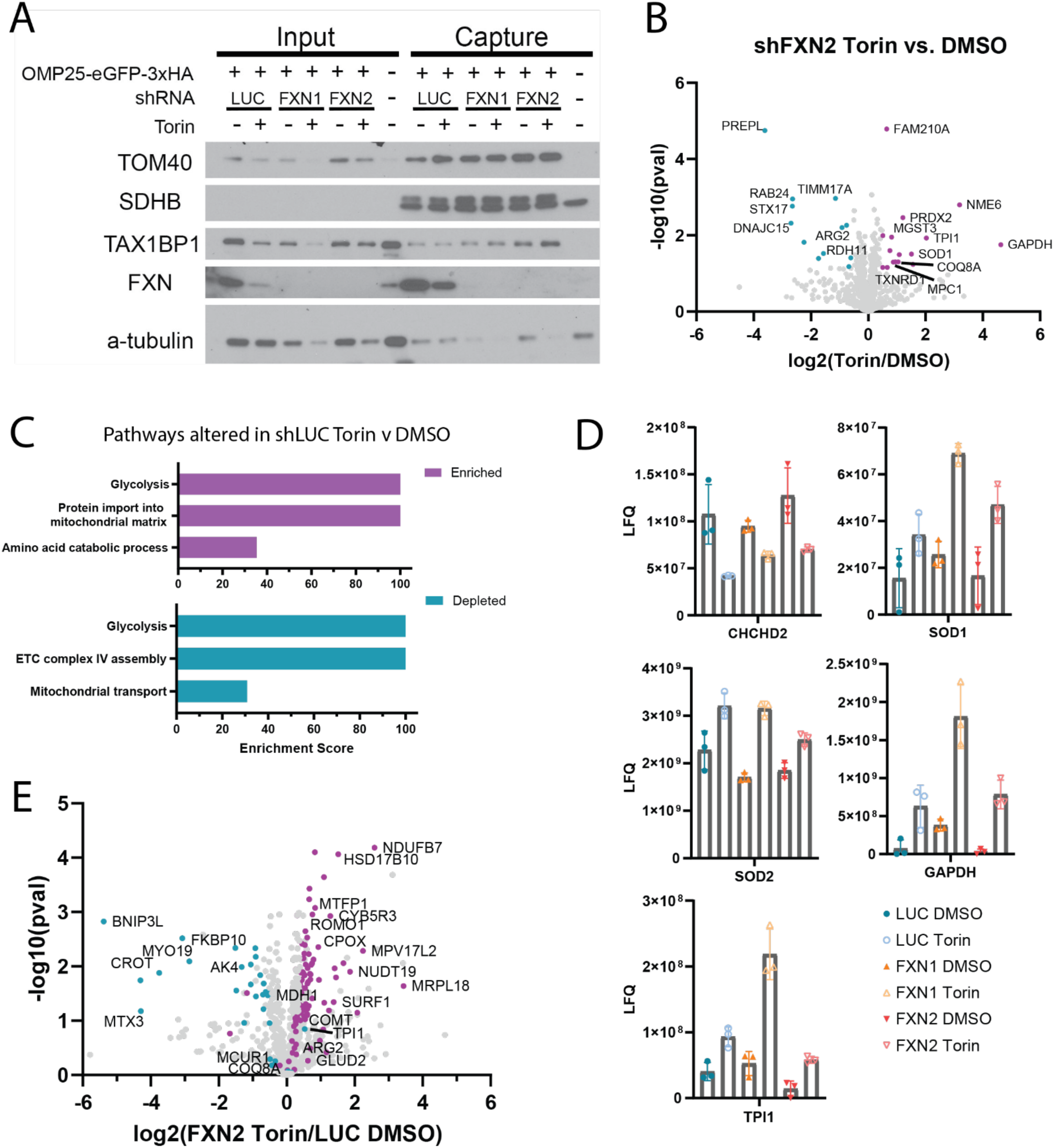
Proteomic analysis of isolated mitochondria give insight into potential mechanisms of mTORC1 control over mitochondrial function. **(A)** Western blot of 10% of each mitoIP proteomic sample prior to normalization to total protein content, demonstrating enrichment of mitochondrial material, loss of cytoplasmic material, and knockdown of FXN **(B)** Volcano plot of mito-IP samples, normalized by total protein content, comparing the mitochondrial proteome of shFXN2 cells in Torin compared to vehicle treatment; **(C)** Enrichment scores for proteins changed during Torin1 treatment in shLUC control cells; **(D)** Bar plots showing quantification levels of glycolysis and stress response genes regulated across all hairpins in Torin treatment; **(E)** Volcano plot comparing the mitochondrial proteome of Torin-treated shFXN2 cells versus control shLUC cells treated with vehicle;

